# Cell entry of SARS-CoV-2 conferred by angiotensin-converting enzyme 2 (ACE2) of different species

**DOI:** 10.1101/2020.06.15.153916

**Authors:** Yan-Dong Tang, Yu-Ming Li, Jing Sun, Hong-Liang Zhang, Tong-Yun Wang, Ming-Xia Sun, Yue-Lin Yang, Xiao-Liang Hu, Jincun Zhao, Xui-Hui Cai

## Abstract

The outbreak of the severe acute respiratory syndrome coronavirus 2 (SARS-CoV-2) poses a huge threat to many countries around the world. However, where is it origin and which animals are sensitive to cross-species transmission is unclear. The interaction of virus and cell receptor is a key determinant of host range for the novel coronavirus. Angiotensin-converting enzyme 2 (ACE2) is demonstrated as the primary entry receptor for SARS-CoV-2. In this study, we evaluated the SARS-CoV-2 entry mediated by ACE2 of 11 different species of animals, and discovered that ACE2 of *Rhinolophus sinicus* (Chinese horseshoe bat), *Felis catus* (domestic cat), *Canis lupus familiaris* (dog), *Sus scrofa* (pig), *Capra hircus* (goat) and especially *Manis javanica* (Malayan pangolin) were able to render SARS-CoV-2 entry in non-susceptible cells. This is the first report that ACE2 of Pangolin could mediate SARS-CoV-2 entry which increases the presume that SARS-CoV-2 may have a pangolin origin. However, none of the ACE2 proteins from *Rhinolophus ferrumequinum* (greater horseshoe bat), *Gallus gallus* (chicken), *Notechis scutatus* (mainland tiger snake), *Mus musculus* (house mouse) rendered SARS-CoV-2 entry. Specifically, a natural isoform of *Macaca mulatta* (Rhesus monkey) ACE2 with a mutation of Y217N was resistance to infection, which rises the possible impact of this type of ACE2 during monkey studies of SARS-CoV-2. Overall, these results clarify that SARS-CoV-2 could engage receptors of multiple species of animals and it is a perplexed work to track SARS-CoV-2 origin and its intermediate hosts.

**IMPORTANCE:** In this study, we illustrated that SARS-CoV-2 is able to engage receptors of multiple species of animals. This indicated that it may be a perplexed work to track SARS-CoV-2 origin and discover its intermediate hosts. This feature of virus is considered to potentiate its diverse cross-species transmissibility. Of note, here is the first report that ACE2 of Pangolin could mediate SARS-CoV-2 entry which increases the possibility that SARS-CoV-2 may have a pangolin origin. And we also demonstrated that not all species of bat were sensitive to SARS-CoV-2 infection. At last, it is also important to detect the expression ratio of the Y217N ACE2 to the prototype in Rhesus monkeys to be recruited for studies on SARS-CoV-2 infection.

## INTRODUCTION

In December 2019, a novel pneumonia, termed as COVID-19 by World Health Organization (WHO) thereafter, emerged in Wuhan, China, and the causative agent was soon identified as a novel coronavirus, which is termed as severe acute respiratory syndrome coronavirus 2 (SARS-CoV-2) by the International Committee on Taxonomy of Viruses, ICTV (1, 2). The SARS-CoV-2 outbreak has been speculatively associated with a seafood market where sales also various land wild animals (3). Bats are recognized as a potential natural reservoir for SARS-CoV-2 (1, 3). However, recently studies indicated that pangolins were also considered as possible natural hosts of this coronavirus (4, 5). Discovering the potential intermediate animal hosts of SARS-CoV-2 and evaluating their possible cross-species transmissibility will be scientifically very important. Unfortunately, we know little about this. Currently, there are no suitable animal models for SARS-CoV-2 infection. A recent study revealed that ferrets and cats were sensitive to SARS-CoV-2 infection, however, these animals showed no clinical symptoms (6). Whether there exist other animal(s) as a candidate SARS-CoV-2 infection model should be further explored.

The interaction between receptor and virus is a key determinant of the host range. It has been demonstrated that SARS-CoV-2 resembles SARS-CoV, which uses angiotensin-converting enzyme 2 (ACE2) as the primary cell entry receptor (1, 7-9). When we retrace the origin of coronavirus, the cell susceptibility to viruses conferred by receptors of speculated animals is a preferential consideration (10, 11). Before clarifying that *Rhinolophus sinicus* is the natural reservoir of SARS-CoV, scientists first evaluated the susceptibility provided by ACE2 from different bat species to SARS-CoV. They found that the ACE2 of *Rhinolophus sinicus* was responsible for the susceptibility to SARS-CoV and subsequently confirmed that *Rhinolophus sinicus* was the natural reservoir of SARS-CoV (10, 12, 13). The Middle East respiratory syndrome coronavirus (MERS-CoV) was also recognized has a bat origin due to that both the MERS-CoV and two MERS-CoV-related viruses from bats could utilize human or bat dipeptidyl peptidase 4 (DPP4) for cell entry (14-16).

Therefore, in this study, we systemically evaluated the ability of SARS-CoV-2 to infect two types of non-susceptible cells utilizing ACE2 proteins from nine different species of animals and the human being to determine its possible origin and further to explore its potentiate cross-species transmission. Our findings provide evidence that SARS-CoV-2 was able to engage broad receptors of different species, which poses a huge challenge to search the animal origin of SARS-CoV-2 for the control and prevention in future.

## RESULT

To investigate which animal’s ACE2 could render SARS-CoV-2 entry, we synthesized the full-length cDNA fragments of ACE2 from 11 species of animals, as well as the human being. These species were *Rhinolophus sinicus* (Chinese horseshoe bat), *Rhinolophus ferrumequinum* (greater horseshoe bat), *Felis catus* (domestic cat), *Capra hircus* (goat), *Canis lupus familiaris* (dog), *Sus scrofa* (pig), *Manis javanica* (Malayan pangolin), *Gallus gallus* (chicken), *Notechis scutatus* (mainland tiger snake), *Mus musculus* (house mouse) and *Macaca mulatta* (Rhesus monkey) and *Homo sapiens* (human). Synthesized DNA fragments were then sub-cloned into the pCAGGS-HA vector for the expression in eukaryotic cells. The origins and GenBank accession numbers of these ACE2 molecules were listed in the Table. We firstly compared the nucleotide sequences of ACE2 coding regions of these animals to that of human. The sequence similarities of these ACE2 cDNAs were exhibited in the Table. Among these sequences, the ACE2 of Rhesus monkey was most close to human, and in contrast, the ACE2 of snake was the farthest. It has been reported that two virus-binding hotspots, Lys31 and Lys353 in hACE2, were critical for SARS-CoV infection (17, 18). In this study, we found that the Lys31 was not conserved in ACE2 of all the 11 animal species observed in this study. However, the Lys353 was conserved in all the 10 animal species except mouse (Table).

Next, we tested whether ACE2 of the nine animal species were able to render SARS-CoV-2 entry to non-susceptible HEK293T cell lines. Different ACE2s could be expressed and presented in the surface of HEK293T cells by IFA (Figure 1). HEK293T Plasmids expressing ACE2 of human and mice were applied as the positive and negative control of the entry assay, respectively. No attempt was made to quantify infection efficiency in this study due to difficulties encountered in conducting experiments under BSL-3 conditions. As expected, the human ACE2 supported SARS-CoV-2 entry whereas mouse ACE2 did not (Figure 1). One previous study indicated that the SARS-CoV outbreak in 17 years ago was originated from *Rhinolophus affinis* (Intermediate horseshoe bat) (12). A recent study further demonstrated that ACE2 of *Rhinolophus sinicus* (Chinese horseshoe bat) also rendered SARS-CoV-2 entry besides SARS-CoV (1). In this study, we was not able to synthesize the ACE2 cDNA of *Rhinolophus affinis* due to the absent of its sequence. Therefore, we synthesized the ACE2 cDNA of *Rhinolophus sinicus* and *Rhinolophus ferrumequinum* (Greater horseshoe bat) to test whether ACE2 of other bat species was responsible for the susceptibility to SARS-CoV-2. Interestingly, we found that the ACE2 of *Rhinolophus ferrumequinum* did not support the SARS-CoV-2 entry as *Rhinolophus sinicus* (Figure 2), suggesting that not all species of bat were sensitive to SARS-CoV-2 infection.

**Figure 1.**
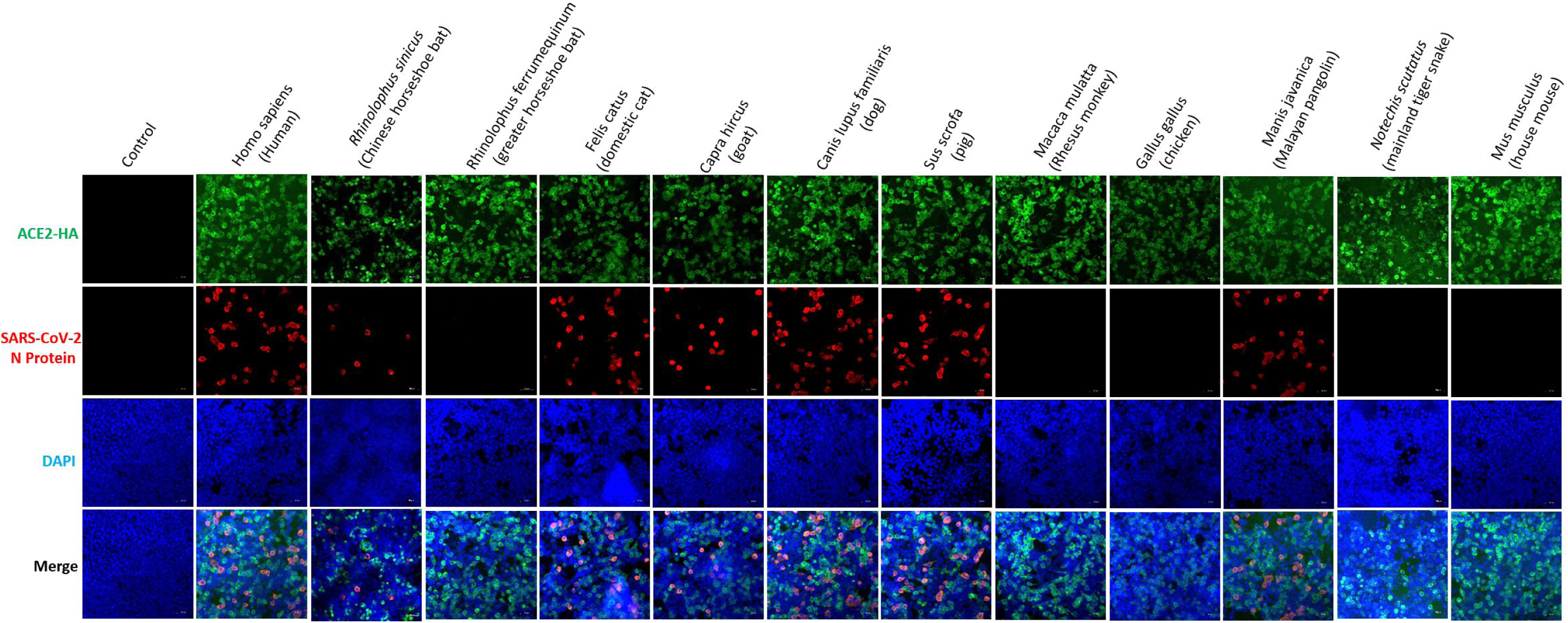
Susceptibility to SARS-CoV-2 of HEK293T cells conferred by different species of ACE2. HEK293T cells were transfected with plasmids expressing indicated ACE2. Cells were infected with 0.5 MOI of SARS-CoV-2 24 h after the transfection, and were detected for the replication of SARS-CoV-2 by IFA.

**Figure 2.**
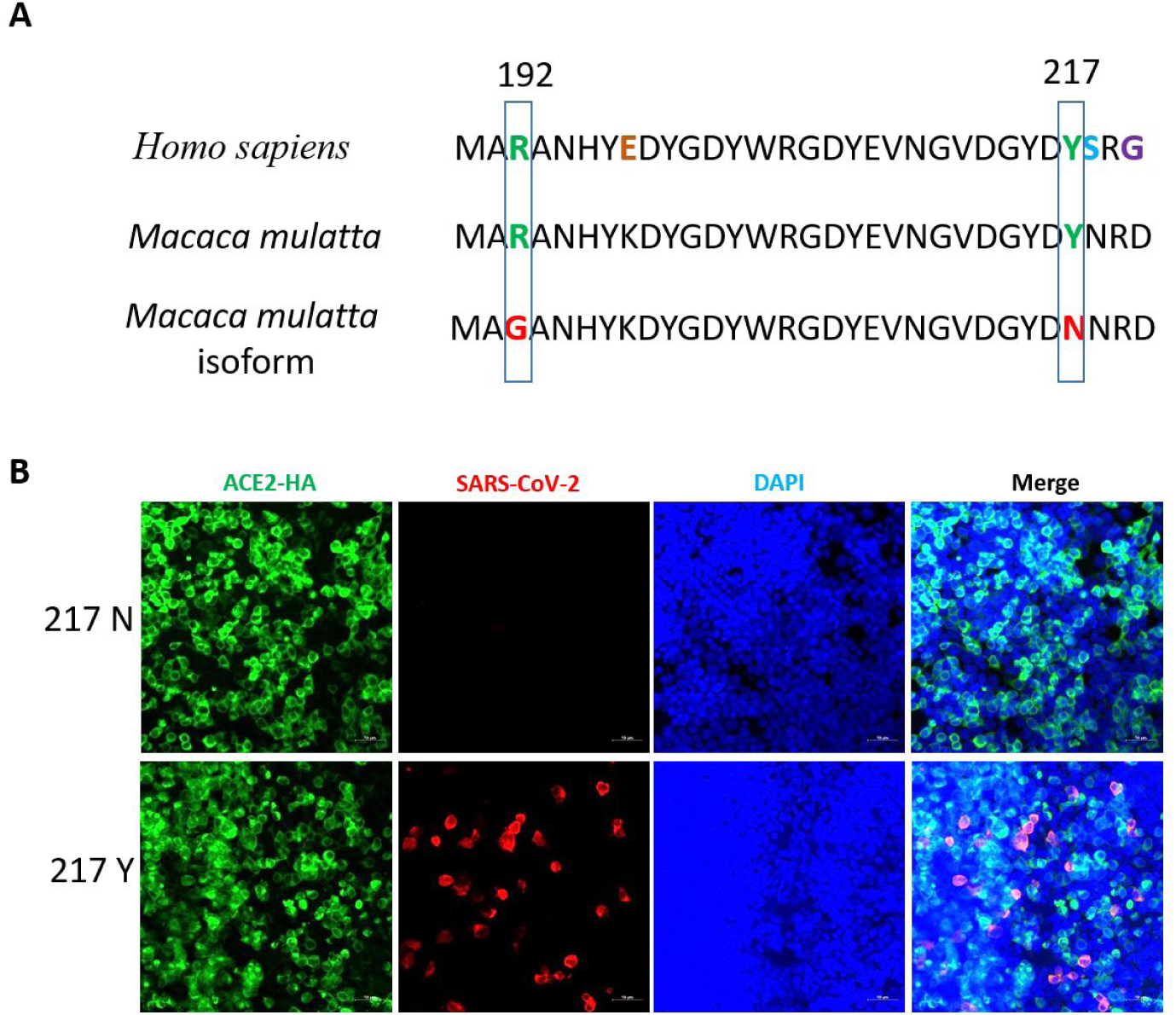
Sequence composition of the Rhesus monkeyACE2 cloned in this study with that of the prototype monkey ACE2 and susceptibility to SARS-CoV-2 of 217 restoration. (**A**)Two sites of natural variation (R192G and Y217N) were identified in the cDNA of Rhesus monkey ACE cloned in this study were compared with the monkey prototype ACE and the human ACE. (**B**) HEK293T cells were transfected with plasmids expressing indicated ACE2. Cells were infected with 0.5 MOI of SARS-CoV-2 24 h after the transfection, and were detected for the replication of SARS-CoV-2 by IFA.

**Table 1.** Nucleotide sequence similarity of various animal ACE-2 to human ACE-2.

| ACE-2 origin | Length of coding sequence (bp) | Similarity to human ACE-2 (%) | Position 31 | Position 353 | GenBank accession number |
| --- | --- | --- | --- | --- | --- |
| *Homo sapiens* (Human) | 2418 | 100 | K | K | AB046569.1 |
| *Rhinolophus ferrumequinum* (Greater horseshoe bat) | 2418 | 86.2 | D | K | AB297479.1 |
| *Rhinolophus sinicus* | 2418 | 85.5 | E | K | GQ262791.1 |
| *Macaca mulatta* (Rhesus monkey)* | 2418 | 96.6 | K | K | NM_001135696.1 |
| *Sus scrofa* (Pig) | 2418 | 84.5 | K | K | NM_001123070.1 |
| *Canis lupus familiaris* (Dog) | 2415 | 87 | K | K | NM_001165260.1 |
| *Capra hircus* (Goat) | 2415 | 85.5 | K | K | KF921008.1 |
| *Felis catus* (Cat) | 2418 | 86.8 | K | K | AY957464.1 |
| *Gallus gallus* (Chicken) | 2427 | 68.1 | E | K | MK560199.1 |
| *Manis javanica* (Malayan pangolin) | 2418 | 86.5 | K | K | XM_017650257.1 |
| *Notechis scutatus* (mainland tiger snake) | 2487 | 66.5 | Q | K | XM_026674969 |
| *Mus musculus* (Mouse) | 2418 | 85.2 | N | H | NM_001130513.1 |

\*NOTE: The sequence of *Macaca mulatta* (Rhesus monkey) here is with a natural mutation Y217N.

A recent study indicated that SARS-CoV-2 did not replicate and shed in dogs, pigs, chickens and ducks, but fairly good in ferrets and effectively in cats (6). Consistently, our study demonstrated that the cat ACE2 supported viral entry (Figure 1). Although pigs, dogs and chickens were non-sensitive to SARS-CoV-2 infection, we know little about its molecular mechanisms and the role of receptor avidity for the resistance. Our data demonstrated that ACE2 proteins of dog and pig supported SARS-CoV-2 entry as that of cat.

Old world monkeys (*Macaca mulatta* and *Macaca fascicularis*) were used as animal models of experimental SARS-CoV-2 infection (19). Surprisingly, we found that the ACE2 of *Macaca mulatta* in our study did not support the SARS-CoV-2 entry as expected (Figure 1). By investigating the monkey ACE2 sequence, we found that an ACE2 isoform of *Macaca mulatta*, which contained two natural variations (R192G and Y217N) comparing with the wild type ACE2, was cloned (Figure 2A). Our data revealed that the ACE2 of *Macaca mulatta* with the Y217N mutation also fault to support SARS-CoV-2 infection. When we restored Y217N mutation as wild type ACE2 of *Macaca mulatta*, N217Y recovery the ability to support SARS-CoV-2 infection (Figure 2B). We noticed that the prototype 217Y was conserved in other species of animals investigated in this study excluding *Macaca mulatta* (data not shown), which suggests that the 217 position is a key residual for SARS-CoV-2 infection.

There is a dispute on if SARS-CoV-2 originated from bats or pangolins (1, 4, 5). It has been demonstrated that a bat ACE2 mediated the SARS-CoV-2 entry (1). However, whether ACE2 of pangolins support the virus entry was unclear. Therefore, we expressed the ACE2 of Malayan pangolins (*Manis javanica*) and tested its role in conferring the susceptibility to SARS-CoV-2. For the first time, we demonstrated that SARS-CoV-2 could engaged the ACE2 of pangolins to entry (Figure 1).

At last, we demonstrated ACE2 of *Notechis scutatus* (mainland tiger snake) could not support SARS-CoV-2 entry as previously predicted (20). And snakes may be not the source of SARS-CoV-2.

## DISSCUSSION

Spike features of coronaviruses and lysosomal proteases of hosts determine the tropism of coronavirus (21). A bat coronavirus RaTG13 in *Rhinolophus affinis* (Intermediate horseshoe bat) from Yunnan exhibits the highest sequence similarity to SARS-CoV-2 until now (1). In this study, we found that the ACE2 of *Rhinolophus ferrumequinum* (Greater horseshoe bat) failed to mediate SARS-CoV-2 entry, whereas ACE2 of *Rhinolophus sinicus* (Chinese horseshoe bat) rendered SARS-CoV-2 entry to non-susceptible cells. In fact, in contrast to the genetically homogenous human ACE2, bat ACE2 proteins have great genetic diversity (13). A number of ACE2 molecules isolated from different bat species failed to mediate efficient SARS-CoV entry (13). A study has reported that *Rhinolophus sinicus* serves as natural reservoirs of SARS-CoV and an isolated bat-origin SARS-CoV-like virus is able to employ ACE2 proteins from humans and civets for cell entry (12). These results suggest that analysis of the receptor-conferred susceptibility to the virus entry is important before investigating for the bat-origin of SARS-CoV-2.

Recently, pangolins are also considered as a possible natural host of SARS-CoV-2 (4, 5). Coronaviruses with high sequence homology were identified in Malayan pangolins (*Manis javanica*) suggesting to be a possible source for the emergence of SARS-CoV-2 (4). We demonstrated that the ACE2 of Malayan pangolin supported SARS-CoV-2 entry in non-susceptible cells (Figure 1). SARS-CoV and MERS-CoV engage receptors of both human and the natural animal hosts (12, 16). Similarly, the ability of pangolin ACE2 to confer the susceptibility to SARS-CoV-2 entry increases the possibility that SARS-CoV-2 originated from pangolins.

In this study, we demonstrated that ACE2 of pig and dog rendered SARS-CoV-2 entry. However, a recent study reported that SARS-CoV-2 replicated poorly in dogs and pigs (6). We speculate that there exist other factors determining the host tropism besides receptor interaction. A recent study demonstrated that pigs and dogs exhibit relatively low levels of ACE2 in the respiratory tract, this may be the reason that SARS-CoV-2 replicated poorly in dogs and pigs (22). Although dogs and pigs may not sensitive for SARS-CoV-2 infection, we do not know whether these animals are appeared as asymptomatic carriers of SARS-CoV-2 in certain environments. In humans asymptomatically infected with SARS-CoV-2, the viral loads were reported similar to that in the symptomatic patients, which implicates the similar risk of viral transmission from asymptomatic carriers (23, 24).

Based on structure analysis of human ACE2 and spikes of SARS-CoV-2, the receptor binding domain (RBD) of the spike takes a more compact conformation than SARS-CoV, implicating a relation to the higher transmission of SARS-CoV-2 than SARS-CoV (25, 26). However, in mouse ACE2, the Lys at site 353 is substituted by His, which does not fit into the virus-receptor binding interface as tightly as the lysine at the same site of human ACE2. Consequently, this may result in the failure of mice ACE2 confer the susceptibility to SARS-CoV-2 entry. A recent publication also demonstrated that the substitution of Lys353 of human ACE2 (hACE2) by Ala was sufficient to abolish the interaction between hACE2 and the S protein of SARS-CoV-2 (27). In addition, although the residue at site 217 of ACE2 is not directly contact to the RBD, this site in Rhesus monkey ACE2 is still critical for SARS-CoV-2 entry. It was observed that the natural variation of Y217N at this site of the monkey ACE-2 significantly reduced the susceptibility to SARS-CoV entry, which demonstrated that this residual variation is responsible for the down regulation of ACE2 expression (28). However, our results showed that this Y217N-isoform of ACE2 expressed at a similar level as the human ACE2 in transfected cells (data not shown). Therefore, the failure of monkey ACE2 isoform to convert the cell susceptibility to SARS-CoV-2 entry is not due to the poor expression of the receptor as previously speculated (28). The detailed mechanism needs further investigation. It is also important to detect the expression ratio of the Y217N ACE2 to the prototype in Rhesus monkeys to be recruited for studies on SARS-CoV-2 infection.

## MATERIALS AND METHODS

### Cells and SARS-CoV-2

HEK293T cells cells were maintained in DMEM (Gibco, USA) with 10% fetal bovine serum (HyClone, USA). The SARS-CoV-2 used in this study (GenBank: MT123290) was isolated from a patient’s throat swab and stored at the Biosafety Level 3 Laboratory of Guangzhou Customs Technical Center.

### Plasmids

The full-length cDNA fragments of different species of ACE2 were synthesized at either the Sangon Biotech (Shanghai, China) or TsingKe Biotech (Nanjing, China). The species and GenBank accession numbers of ACE2 sequences were listed in the Table. Synthesized DNA fragments were then sub-cloned into a eukaryotic expression vector pCAGGS-HA for the expression in human cell lines.

### Sequence analysis

ACE2 sequences of 12 different species were acquired from NCBI and their alignment were assessed using the ClustalW method in Lasergene software (Version 7.1) (DNASTAR Inc., USA).

### Entry assay

HEK293T cells were plated in 48-well plates, and were transfected with indicated plasmids by the X-tremeGENE HP DNA Transfection Reagent (Roche, USA) when the cell confluence was up to 90%. Cells were infected with 0.5 MOI of SARS-CoV-2 24 h after being transfected. The detection of infected cells were performed 12 h late by using an immunofluorescence assay as described previously (29). An HA-Alexa Fluor 488 monoclonal antibody (Thermo Fisher Scientific, USA) was used to stain ACE2 with an HA tag. The nucleoprotein (N) of SARS-CoV was detected for infection and replication of the virus using an N-specific polyclonal antibody (Sinobiological, China), and a donkey anti-rabbit IgG (H+L) labeled with Cy3 (Jacksion, USA) was used as the secondary antibody. All the cells were stained with DAPI (Sigma, USA) for nuclear visualization.

## Compliance with ethical standards

### Funding

This study was funded by grants from The National Key Research and Development Program of China (2018YFC1200100, 2018ZX10301403), the emergency grants for prevention and control of SARS-CoV-2 of Ministry of Science and Technology (2020YFC0841400) and Guangdong province (2020B111108001, 2018B020207013).

### Ethical approval

This article does not contain any studies with human participants

## or Acknowledgments

The authors are grateful to Prof. Jian-Hua Zhou for kindly revising the manuscript.

## Reference

1. Zhou P, Yang XL, Wang XG, Hu B, Zhang L, Zhang W, Si HR, Zhu Y, Li B, Huang CL, Chen HD, Chen J, Luo Y, Guo H, Jiang RD, Liu MQ, Chen Y, Shen XR, Wang X, Zheng XS, Zhao K, Chen QJ, Deng F, Liu LL, Yan B, Zhan FX, Wang YY, Xiao GF, Shi ZL. 2020. A pneumonia outbreak associated with a new coronavirus of probable bat origin. Nature 579:270–273.

2. Wu F, Zhao S, Yu B, Chen YM, Wang W, Song ZG, Hu Y, Tao ZW, Tian JH, Pei YY, Yuan ML, Zhang YL, Dai FH, Liu Y, Wang QM, Zheng JJ, Xu L, Holmes EC, Zhang YZ. 2020. A new coronavirus associated with human respiratory disease in China. Nature 579:265–269.

3. Lu R, Zhao X, Li J, Niu P, Yang B, Wu H, Wang W, Song H, Huang B, Zhu N, Bi Y, Ma X, Zhan F, Wang L, Hu T, Zhou H, Hu Z, Zhou W, Zhao L, Chen J, Meng Y, Wang J, Lin Y, Yuan J, Xie Z, Ma J, Liu WJ, Wang D, Xu W, Holmes EC, Gao GF, Wu G, Chen W, Shi W, Tan W. 2020. Genomic characterisation and epidemiology of 2019 novel coronavirus: implications for virus origins and receptor binding. Lancet 395:565–574.

4. Lam TT, Shum MH, Zhu HC, Tong YG, Ni XB, Liao YS, Wei W, Cheung WY, Li WJ, Li LF, Leung GM, Holmes EC, Hu YL, Guan Y. 2020. Identifying SARS-CoV-2 related coronaviruses in Malayan pangolins. Nature doi: 10.1038/s41586-020-2169-0.

5. Zhang T, Wu Q, Zhang Z. 2020. Probable Pangolin Origin of SARS-CoV-2 Associated with the COVID-19 Outbreak. Curr Biol doi: 10.1016/j.cub.2020.03.022.

6. Shi J, Wen Z, Zhong G, Yang H, Wang C, Huang B, Liu R, He X, Shuai L, Sun Z, Zhao Y, Liu P, Liang L, Cui P, Wang J, Zhang X, Guan Y, Tan W, Wu G, Chen H, Bu Z. 2020. Susceptibility of ferrets, cats, dogs, and other domesticated animals to SARS-coronavirus 2. Science doi: 10.1126/science.abb7015.

7. Li W, Moore MJ, Vasilieva N, Sui J, Wong SK, Berne MA, Somasundaran M, Sullivan JL, Luzuriaga K, Greenough TC, Choe H, Farzan M. 2003. Angiotensin-converting enzyme 2 is a functional receptor for the SARS coronavirus. Nature 426:450–4.

8. Hoffmann M, Kleine-Weber H, Schroeder S, Kruger N, Herrler T, Erichsen S, Schiergens TS, Herrler G, Wu NH, Nitsche A, Muller MA, Drosten C, Pohlmann S. 2020. SARS-CoV-2 Cell Entry Depends on ACE2 and TMPRSS2 and Is Blocked by a Clinically Proven Protease Inhibitor. Cell doi: 10.1016/j.cell.2020.02.052.

9. Wan Y, Shang J, Graham R, Baric RS, Li F. 2020. Receptor Recognition by the Novel Coronavirus from Wuhan: an Analysis Based on Decade-Long Structural Studies of SARS Coronavirus. J Virol 94.

10. Li W, Wong SK, Li F, Kuhn JH, Huang IC, Choe H, Farzan M. 2006. Animal origins of the severe acute respiratory syndrome coronavirus: insight from ACE2-S-protein interactions. J Virol 80:4211–9.

11. Ren W, Qu X, Li W, Han Z, Yu M, Zhou P, Zhang SY, Wang LF, Deng H, Shi Z. 2008. Difference in receptor usage between severe acute respiratory syndrome (SARS) coronavirus and SARS-like coronavirus of bat origin. J Virol 82:1899–907.

12. Ge XY, Li JL, Yang XL, Chmura AA, Zhu G, Epstein JH, Mazet JK, Hu B, Zhang W, Peng C, Zhang YJ, Luo CM, Tan B, Wang N, Zhu Y, Crameri G, Zhang SY, Wang LF, Daszak P, Shi ZL. 2013. Isolation and characterization of a bat SARS-like coronavirus that uses the ACE2 receptor. Nature 503:535–8.

13. Hou Y, Peng C, Yu M, Li Y, Han Z, Li F, Wang LF, Shi Z. 2010. Angiotensin-converting enzyme 2 (ACE2) proteins of different bat species confer variable susceptibility to SARS-CoV entry. Arch Virol 155:1563–9.

14. Raj VS, Mou H, Smits SL, Dekkers DH, Muller MA, Dijkman R, Muth D, Demmers JA, Zaki A, Fouchier RA, Thiel V, Drosten C, Rottier PJ, Osterhaus AD, Bosch BJ, Haagmans BL. 2013. Dipeptidyl peptidase 4 is a functional receptor for the emerging human coronavirus-EMC. Nature 495:251–4.

15. Lau SKP, Fan RYY, Luk HKH, Zhu L, Fung J, Li KSM, Wong EYM, Ahmed SS, Chan JFW, Kok RKH, Chan KH, Wernery U, Yuen KY, Woo PCY. 2018. Replication of MERS and SARS coronaviruses in bat cells offers insights to their ancestral origins. Emerg Microbes Infect 7:209.

16. Yang Y, Du L, Liu C, Wang L, Ma C, Tang J, Baric RS, Jiang S, Li F. 2014. Receptor usage and cell entry of bat coronavirus HKU4 provide insight into bat-to-human transmission of MERS coronavirus. Proc Natl Acad Sci U S A 111:12516–21.

17. Li F. 2008. Structural analysis of major species barriers between humans and palm civets for severe acute respiratory syndrome coronavirus infections. J Virol 82:6984–91.

18. Wu K, Peng G, Wilken M, Geraghty RJ, Li F. 2012. Mechanisms of host receptor adaptation by severe acute respiratory syndrome coronavirus. J Biol Chem 287:8904–11.

19. Lu S, Zhao, Y., Yu, W., Yang, Y., Gao, J., Wang, J., Kuang, D., Yang, M., Yang, J., Ma, C., et al. 2020. Comparison of SARS-CoV-2 infections among 3 species of non-human primates. bioRxiv.

20. Ji W, Wang W, Zhao X, Zai J, Li X. 2020. Cross-species transmission of the newly identified coronavirus 2019-nCoV. J Med Virol 92:433–440.

21. Zheng Y, Shang J, Yang Y, Liu C, Wan Y, Geng Q, Wang M, Baric R, Li F. 2018. Lysosomal Proteases Are a Determinant of Coronavirus Tropism. J Virol 92.

22. Zhai X, Sun J, Yan Z, Zhang J, Zhao J, Zhao Z, Gao Q, He WT, Veit M, Su S. 2020. Comparison of SARS-CoV-2 spike protein binding to ACE2 receptors from human, pets, farm animals, and putative intermediate hosts. J Virol doi: 10.1128/JVI.00831-20.

23. Zou L, Ruan F, Huang M, Liang L, Huang H, Hong Z, Yu J, Kang M, Song Y, Xia J, Guo Q, Song T, He J, Yen HL, Peiris M, Wu J. 2020. SARS-CoV-2 Viral Load in Upper Respiratory Specimens of Infected Patients. N Engl J Med 382:1177–1179.

24. Rothe C, Schunk M, Sothmann P, Bretzel G, Froeschl G, Wallrauch C, Zimmer T, Thiel V, Janke C, Guggemos W, Seilmaier M, Drosten C, Vollmar P, Zwirglmaier K, Zange S, Wolfel R, Hoelscher M. 2020. Transmission of 2019-nCoV Infection from an Asymptomatic Contact in Germany. N Engl J Med 382:970–971.

25. Shang J, Ye G, Shi K, Wan Y, Luo C, Aihara H, Geng Q, Auerbach A, Li F. 2020. Structural basis of receptor recognition by SARS-CoV-2. Nature doi: 10.1038/s41586-020-2179-y.

26. Yan R, Zhang Y, Li Y, Xia L, Guo Y, Zhou Q. 2020. Structural basis for the recognition of the SARS-CoV-2 by full-length human ACE2. Science doi: 10.1126/science.abb2762.

27. Wang Q, Zhang Y, Wu L, Niu S, Song C, Zhang Z, Lu G, Qiao C, Hu Y, Yuen KY, Zhou H, Yan J, Qi J. 2020. Structural and Functional Basis of SARS-CoV-2 Entry by Using Human ACE2. Cell doi: 10.1016/j.cell.2020.03.045.

28. Chen Y, Liu L, Wei Q, Zhu H, Jiang H, Tu X, Qin C, Chen Z. 2008. Rhesus angiotensin converting enzyme 2 supports entry of severe acute respiratory syndrome coronavirus in Chinese macaques. Virology 381:89–97.

29. Yang YL, Meng F, Qin P, Herrler G, Huang YW, Tang YD. 2020. Trypsin promotes porcine deltacoronavirus mediating cell-to-cell fusion in a cell type-dependent manner. Emerg Microbes Infect 9:457–468.

